# Sex differences in cardiomyocyte proteomic responses to the cardiotoxic chemotherapy drug doxorubicin

**DOI:** 10.64898/2026.04.21.715711

**Authors:** Sanela Dozic, Michael G. Leeming, Keshava K. Datta, Hitesh Kore, Erin J. Howden, Benjamin L. Parker, Lea M.D. Delbridge, Kate L. Weeks

## Abstract

Anthracyclines, including doxorubicin, are widely used chemotherapeutic agents, but dose-dependent cardiotoxicity limits their clinical utility and increases the risk of heart failure in cancer survivors. In paediatric patients, female sex is a significant risk factor for anthracycline-associated cardiotoxicity, yet pre-clinical studies rarely investigate sex differences in immature hearts. Here, we provide a proteomic dataset from primary cardiomyocytes, isolated from postnatal day 2 rat hearts and treated with a clinically relevant concentration of doxorubicin. Analysis of proteins present in all samples identified candidates previously shown to be regulated by doxorubicin in adult hearts, as well as candidates that may be specifically regulated in young hearts. This dataset provides a resource for generating hypotheses on molecular mechanisms contributing to sex differences in juvenile doxorubicin-induced cardiotoxicity.

## Background & Summary

Anthracyclines are highly effective chemotherapeutic agents used to treat a range of solid tumours and haematological malignancies in both adults and children.^1–3^ However, the clinical utility of anthracyclines is constrained by dose-dependent cardiotoxicity, which can lead to lifelong morbidity and an increased risk of heart failure.^4^ Age and sex are established risk factors for anthracycline-associated cardiotoxicity, with female paediatric cancer survivors at higher risk of developing congestive heart failure, and more likely to die from cardiac diseases, than age-matched male cancer survivors.^5,6^ Interestingly, this sex-based risk profile changes for adult cancer patients, likely due to the cardioprotective effects of estrogen for women.^7^ Elucidating the mechanisms by which anthracyclines damage the myocardium in female and male patients at different life stages will inform the development of gender-optimised therapeutic strategies to mitigate cardiac injury and dysfunction resulting from chemotherapy.

Mechanistic studies investigating the molecular basis for anthracycline-associated cardiotoxicity have primarily used adult rodents, and support anecdotal evidence in adult cancer patients that female sex is cardioprotective.^8^ Very few studies have modelled juvenile anthracycline-associated cardiotoxicity, with most not considering sex in the experimental design. Several developmental and physiological differences in rodents make it challenging to model the human pre-pubertal period, particularly for multi-week drug treatment regimes. Weaning of rodents typically occurs from postnatal day 21, with reports of sexual maturity reached by 4 weeks of age.^9^ Studies modelling anthracycline-associated cardiotoxicity in young animals typically administer doxorubicin (DOX) daily for 7-14 days^10,11^ or weekly for 3-7 weeks^12–15^, commencing at 2-7 weeks of age. Thus, many published models of juvenile anthracycline-associated cardiotoxicity likely reflect responses of young adult animals under the influence of sex hormones.

Here, we make available a dataset which allows comparison of the male and female cardiomyocyte proteome, in the absence and presence of DOX. We used primary cardiomyocytes isolated from postnatal day 2 rat pups, which are widely used to investigate cardiac physiology and signalling^16–19^ and have previously been shown to recapitulate gene expression patterns of paediatric dilated cardiomyopathy.^20,21^ A novel aspect of our experimental design was to isolate cardiomyocytes from individual hearts instead of pooling cardiomyocytes from multiple hearts, allowing paired (i.e. within-animal) analysis of DOX- and vehicle-treated cells. This *in vitro* approach provides an opportunity to explore mechanisms and develop hypotheses regarding DOX-induced cardiotoxicity, and to provide insight into sex differences of young hearts exposed to cardiotoxic chemotherapeutic agents.

## Methods

### Experimental animals & ethics approval

All animal experiments were performed according to the guidelines outlined in the Australian Code of Practice for the Care and Use of Animals for Scientific Purposes (8^th^ edition, updated 2021). Experimental protocols were approved by the University of Melbourne Animal Ethics Committee. Sprague Dawley rat pups aged 2-days old were obtained from the University of Melbourne Biological Research Facility for isolation of neonatal rat ventricular myocytes (NRVM).

### Cardiomyocyte isolation

NRVM were isolated in two independent batches, each comprising of three male and three female rat hearts (*n*=6 per batch; *n*=12 in total). Ventricles were cut into smaller pieces (1-2 mm^3^) and transferred into gentleMACS C Tubes (Miltenyi Biotec, #130-093-237), with one heart allocated per tube. The Neonatal Heart Dissociation Kit (Miltenyi Biotec, #130-098-373) was used to enzymatically digest ventricle tissue. Briefly, 2.5 mL of enzyme master mix was made up from 2362.5 µL of enzyme mix 1 (62.5 µL enzyme P, 2300 µL buffer X) and 137.5 µL of enzyme mix 2 (25 µL buffer Y, 12.5 µL enzyme A, 100 µL enzyme D). Each C Tube containing ventricle tissue and 2.5 mL enzyme master mix was placed onto the gentleMACS Octo Dissociator with heaters (Miltenyi Biotec, #130-096-427) and digested using the 37C_mr_NHDK_1 program. After termination of the program, C Tubes were detached and the enzymatic reaction terminated with the addition of 7.5 mL cold 10% newborn calf serum (NBCS; ThermoFisher, #16010159) prepared in Dulbecco’s Modified Eagle Medium (DMEM; ThermoFisher, #11966025) containing 5 mM glucose (Sigma, #RDD016) and 1% antibiotic-antimycotic (ThermoFisher, #15240062). The C tubes were briefly spun at 100 x g for 1 minute to maximise sample transfer. Cells were filtered through a 70 µm cell strainer (Rowe Scientific, #PC0349) and centrifuged at 600 x g for 5 minutes. The cell pellets were resuspended with 11 mL 10%NBCS and plated in culture dishes for 45-60 minutes (37 °C, 5% CO_2_). This was repeated a second time to facilitate fibroblast adhesion, thereby enriching the cardiomyocyte population. Isolated cells were treated with 1% bromodeoxyuridine (BrdU; Sigma, #B5002) to limit non-myocyte proliferation and plated at 500,000 cells/well of a 12 well plate. After a 48-hour incubation in 10%NBCS (37°C, 5% CO_2_), media was replaced with serum-free DMEM for 16-18 hours to achieve cell cycle synchronisation.

### DOX treatment

After overnight incubation with DMEM, the media was replaced with 10%NBCS. NRVM were then treated with doxorubicin (DOX; Cayman, #15007) at a final concentration of 0.3 µM for 24 hours. Control NRVM were treated with an equivalent volume of saline.

### Protein extraction and normalisation

NRVM were washed with phosphate-buffered saline (PBS; ThermoFisher, #10010023) and lysed in 100 µL Radioimmunoprecipitation Assay (RIPA) lysis and extraction buffer (ThermoFisher, #89901), containing phosphatase (Sigma, #4906845001) and protease (Sigma, #11836170001) inhibitors. Cells were scraped and transferred to tubes for heat denaturation at 95 °C for 5 minutes. Samples were centrifuged at 14,000 x g for 15 minutes (4 °C) to remove insoluble debris and the supernatant transferred to fresh tubes for ultrafiltration. Spin filters (Amicon Ultra-0.5 mL, #UFC501096) were used to concentrate the protein component into ∼30 µL volumes. Briefly, the samples were centrifuged at 14,000 x g for 15 minutes. The solute concentrate was recovered by inverting spin filters into fresh tubes and centrifuging the samples at 1,000 x g for 2 minutes. Samples were made up to 35 µL volumes with MilliQ-H_2_O, with 5 µL aliquoted to determine protein lysate concentration using a microBCA assay (ThermoFisher, #23235). 25 µg protein equivalent lysates were utilised to prepare samples for proteomics.

### Sample preparation for proteomics

Filter Aided Sample Preparation (FASP) was performed on 25 µg cell lysates, made up to 30 µL with RIPA lysis and extraction buffer. Samples were reduced with 3 µL of 10 mM TCEP (45 minutes at 55 °C). Proteins were denatured with addition of 200 µL Urea Tris-HCl solution and centrifugation (14,000 x g for 15 minutes). Alkylation was achieved with addition of 100 µL of 5 mM Iodoacetamide (600 rpm for 1 minute), followed by incubation in the dark for 20 minutes and centrifugation (14,000 x g for 15 minutes). Two iterations were performed with addition of 100 µL of 8 M Urea Tris-HCl (14,000 x g for 15 minutes) and 100 µL of 50 mM ammonium bicarbonate (14,000 x g for 10 minutes). For tryptic digestion, 75 µL of trypsin solution (0.003 µg/µL) was added (600 rpm for 1 minute) and samples incubated overnight (37°C for 15-18 hours). Two iterations of 40 µL of 50 mM ammonium bicarbonate were added (14,000 x g for 10 minutes). Peptides were washed with 50 µL of 0.5 M sodium chloride (14,000 x g for 10 minutes). Acidification was achieved with addition of 10% formic acid to a final pH of 2-3. An Oasis HLB 96 well plate and vacuum manifold was used for de-salting. Briefly, wells were activated twice with 400 µL of 100% LC-MS grade methanol. The HLB plate was equilibrated with two additions of 400 µL of 0.1% formic acid. Samples were dispensed into the HLB plate and washed three times with 400 µL 0.1% formic acid prior to elution with 2 × 75 µL of 50% acetonitrile/0.1% formic acid. Sample were collected into LoBind tubes and vacuum dried with liquid nitrogen for ∼3.5 hours and stored at -80 °C until required. Pellets were resuspended in 30 µL of 0.1% formic acid and sonicated for 5 min (10 sec on 10 sec off) at 70% amplitude at 4 °C. A microBCA assay was performed to determine the final concentration, with samples diluted to 0.1 µg/µL with 0.1% formic acid. Finally, samples were centrifuged (14,000 x g for 10 minutes at 4 °C) and 20 µL volumes aliquoted for mass spectrometric analysis.

### LC-MS/MS data acquisition

An Orbitrap Exploris 480 mass spectrometer (ThermoFisher Scientific, San Jose, USA) interfaced with an Ultimate-3000 UHPLC (ThermoFisher Scientific, San Jose, USA) was utilised to acquire LC-MS/MS data. Peptides were first loaded onto an Acclaim™ PepMap™ C18 trap column (ThermoFisher Scientific, Bremen, Germany) at 5µL/min and washed for 6 minutes. The trap was then brought in-line with the analytical column which was a 50 cm PepMap™ C18 column (ThermoFisher Scientific, Bremen, Germany). The peptides were resolved at a flow rate of 300 nL/min. The solvent system consisted of solvent A (0.1% formic acid, 5% DMSO, in water) and solvent B (0.1% formic acid, 5% DMSO in acetonitrile). The gradient consisted of a linear increase of solvent B from 3% to 25% over 76 minutes and ramp up of 25% to 40% over 4 minutes and 40% to 80% over a minute. The flow was held at 80% B for 3 minutes to wash the column. Flow was dropped to 3% B and the column was equilibrated for 5 minutes. The total runtime was 95 minutes. The mass spectrometer was operated on a Data Independent Mode. The instrument settings were: MS1 Resolution-120,000; Mass Range-350–1400 *m/z*; Normalised AGC Target-250%, Maximum injection time 50 ms. MS2: Isolation window-13.7 *m/z* leading to 50 scan events; Normalised HCD collision energy 30%; Normalised AGC target 2000%, Maximum injection time 55 ms.

### Data processing & analysis

Mass spectrometry data was processed with FragPipe (v.23) using label free quantification-match between runs (LFQ-MBR). A *rattus norvegicus* (UP000234681) protein sequence database was obtained from UniProt, containing 39,538 entries (640 Swiss-Prot and 38,898 TrEMBL), with subsequent addition of decoys and contaminants. Precursor and fragment mass tolerance was set to 20 ppm for MSFragger analysis, with mass calibration and parameter optimisation enabled. For enzymatic cleavage, “stricttrypsin” was selected, with peptide length and mass set to 7-50 and 500-5,000 Da, respectively. Resultant MaxLFQ intensities were used for differential protein abundance analysis using the limma R package. Proteins were filtered to include only those present in all samples. Log_2_ transformed, quantile normalised intensities were analysed using a linear model to include treatment, with sex and batch as covariates. Bioreplicate blocking was applied to estimate correlation between paired samples. Moderated t-statistics were computed by linear model fitting, group contrasts and empirical Bayes moderation.

### Data Records

The mass spectrometry proteomics data have been deposited to the ProteomeXchange Consortium (http://proteomecentral.proteomexchange.org) via the PRIDE partner repository^22^ with the dataset identifier PXD076454. The raw data files carry the project identifier ‘250902_SD-11625’ tagged with the respective sample number (e.g. _01). Sample details are provided in the ‘Experimental_Annotation’ file. Raw data files were processed with FragPipe and the resulting ‘combined_protein’ file used for limma analysis.

### Technical Validation

#### Cell isolations & sample preparation

Ventricular myocytes were mechanically and enzymatically isolated from individual rat pup hearts using an Octo Dissociator, which enabled isolation of cells from multiple hearts in parallel (Fig 1). Two batches of six hearts were isolated on separate days to minimise variability in the length of time that hearts were exposed to the enzyme mix during the isolation procedure. To control for order effects, sex was alternated within each batch (Batch 1: M-F-M-F-M-F, Batch 2: F-M-F-M-F-M). Cardiomyocytes from each heart were divided across two wells in a 12-well plate, for treatment with vehicle and DOX, with each plate including at least one male and one female heart to control for plate effects. Gross morphology of vehicle-treated NRVM was normal, with fewer adherent cells in DOX-treated wells after 24 hours (Fig 2A-B). Importantly, despite a small isolation order effect on protein yield (Fig 2C), there were no differences in total protein between male and female samples (Fig 2D). DOX treatment reduced protein yield in both sexes (Fig 2D), consistent with our observation of fewer adherent cells by microscopy (Fig 2A-B). Protein amount was normalised prior to FASP, and equal amounts of peptide were used for mass spectrometry.

**Figure 1:**
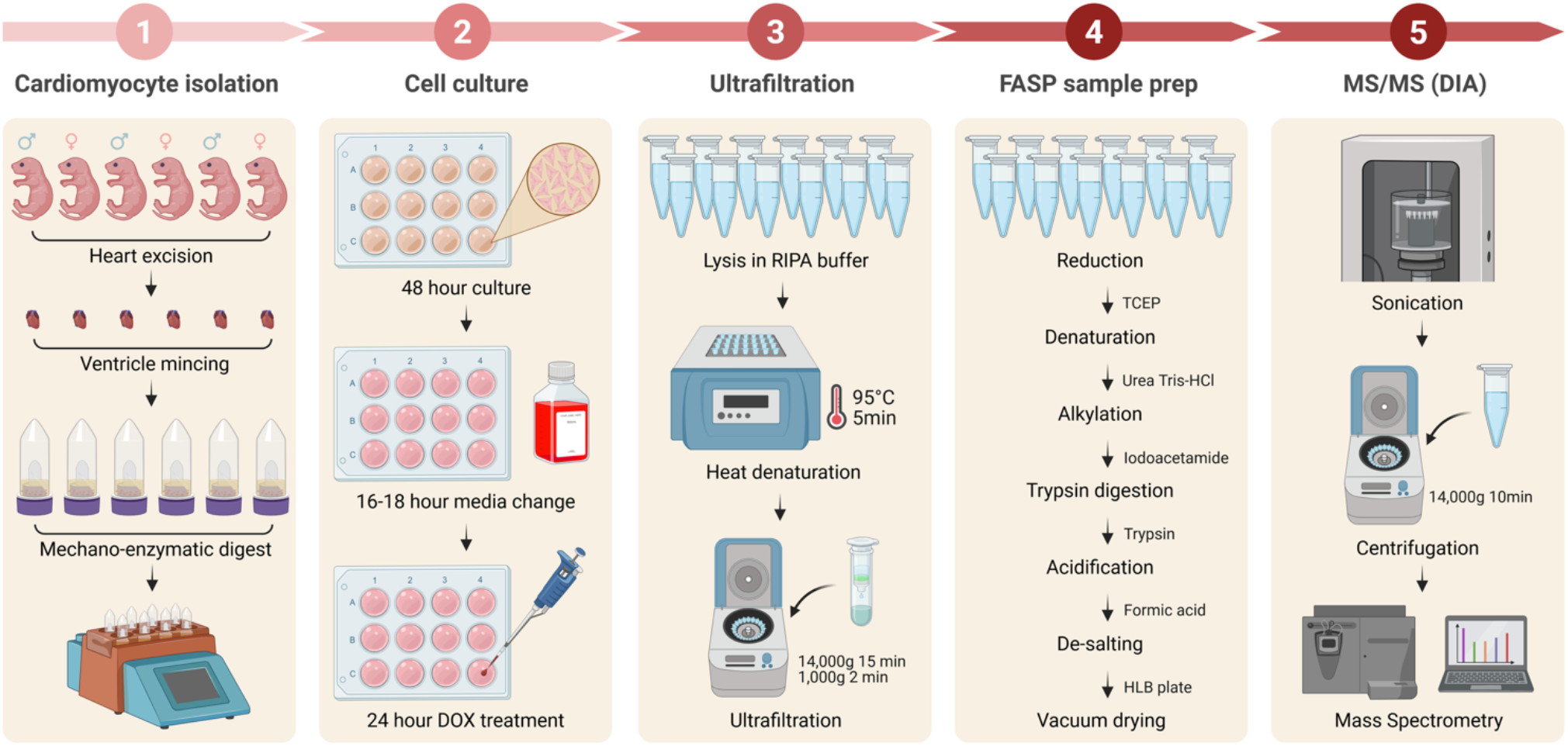
Experimental design and workflow for proteomic data analysis of male and female cardiomyocytes treated with doxorubicin. Individual hearts were excised from postnatal day 2 rat pups and ventricular tissue digested using an Octo Dissociator and Kit. Neonatal rat ventricular myocytes were cultured in 10% newborn calf serum for 48 hours prior to overnight serum depletion (16-18 hours). Cultured myocytes were treated with vehicle and 0.3 μM doxorubicin (Dox) for 24 hours. Filter-aided sample preparation (FASP) was performed on 25 µg cell lysates for data-independent acquisition (DIA) mass spectrometry. Image created with Biorender.

**Figure 2:**
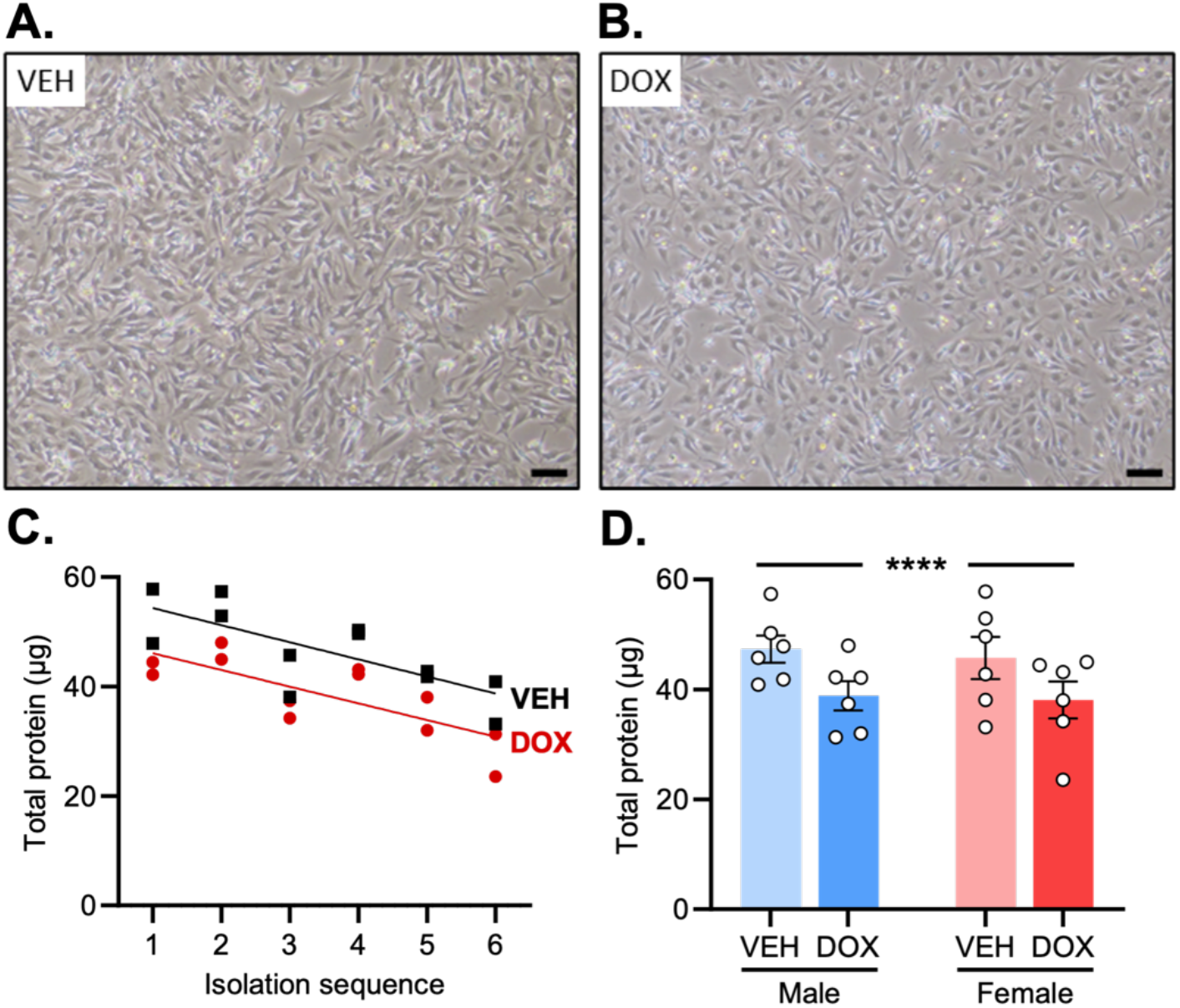
Sample preparation for proteomics workflow. Representative microscopy images of neonatal rat ventricular myocytes cultured in 10% newborn calf serum and treated with a) vehicle and b) 0.3 μM doxorubicin (Dox) for 24 hours. Total protein quantified and grouped by c) isolation sequence (*n*=2, batch) and d) sex and treatment. Data presented as mean ± SEM, with two-way RM ANOVA for statistical testing. \*\*\*\**p*<0.0001, treatment effect. Scale bar: 100 µm.

#### Quality control and main DOX effect

Mass spectra were obtained from 22 samples, with two samples (Dox_5 and Vehicle_9) excluded from further analysis due to comparably lower protein counts (Fig 3A). A threshold was applied to only include proteins detected in all samples. The resulting 2,229 proteins quantified were comparable in distribution following log_2_ transformation & quantile normalisation (Fig 3B). The median percentage coefficient of variation was ∼20%, except for the Male DOX group which showed greater variability due to the smaller sample size (*n*=4; Fig 3C). There was a high degree of correlation between all samples (Pearson correlation coefficient 0.88-0.98; Fig 3D). Analysis of quantitative data with limma identified 140 proteins that were differentially abundant between conditions (Fig 4A). STRING analysis with Gene Ontology enrichment identified biological processes that have previously been associated with DOX cardiotoxicity,^23–27^ including downregulation of proteins involved in translation, actin cytoskeleton organization and biosynthetic processes, and upregulation of proteins involved in the cellular response to a chemical stimulus, response to a toxic substance, and metabolic processes (Fig 4B, C). The top 50 significantly differentially abundant proteins (Fig 4D) included proteins with established roles in DOX-induced cardiac pathology (e.g. calponin 1, CNN1)^28^ and proteins modulated in human heart tissue exposed to DOX (e.g. heat shock protein family A member 5, HSPA5)^29^.

**Figure 3:**
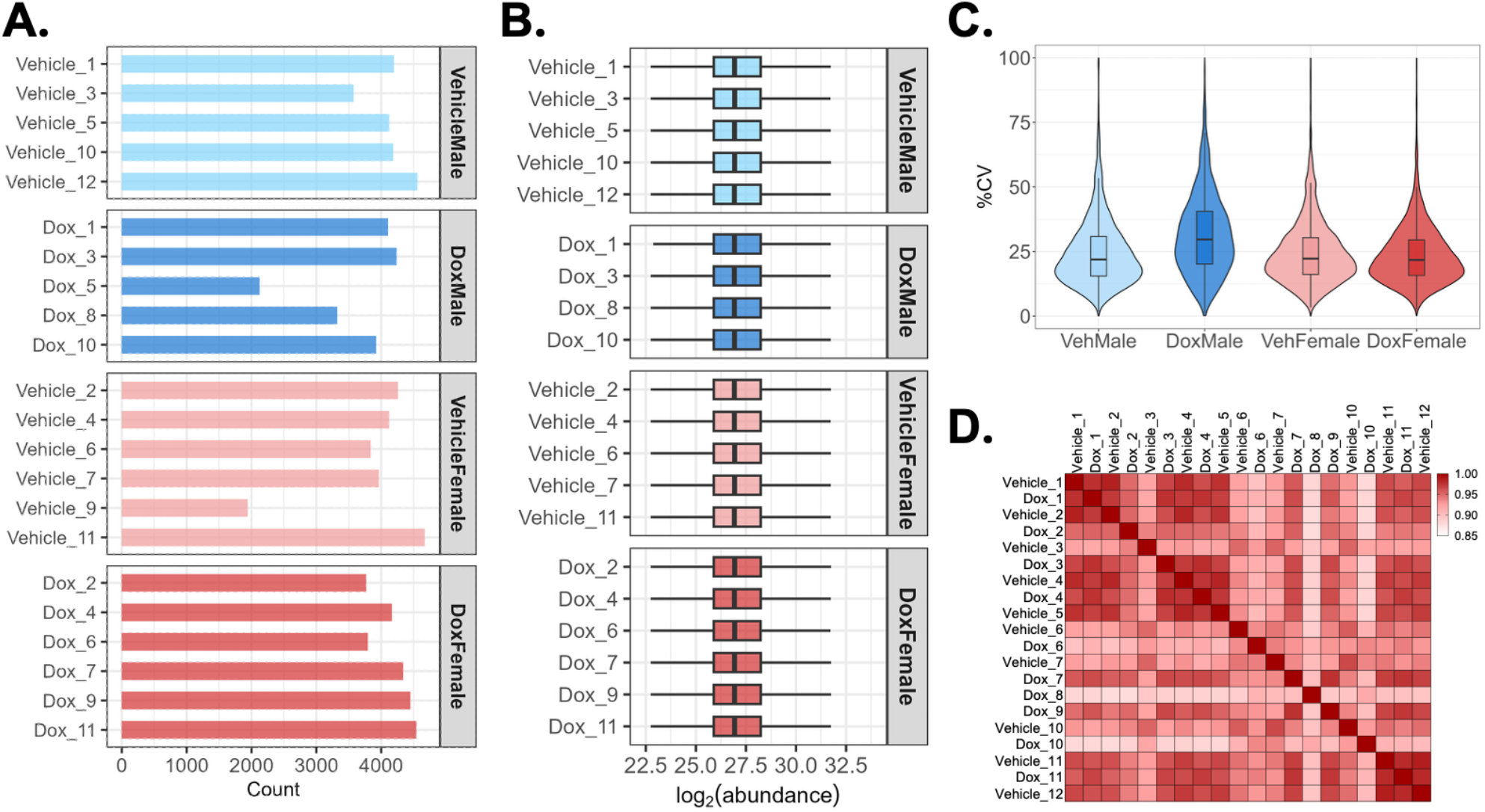
Quality control of the cardiomyocyte proteome in response to doxorubicin. Neonatal rat ventricular myocytes cultured in 10% newborn calf serum and treated with vehicle and 0.3 μM doxorubicin (Dox) for 24 hours. a) Number of identified proteins per sample (*n*=22), with Dox_5 and Vehicle_9 excluded from subsequent analyses. b) Protein abundance was log_2_ transformed and quantile normalised. c) Coefficient of variance was calculated by back-transforming protein abundance to the linear scale. d) Heatmap of Pearson correlation coefficients between the 20 samples.

**Figure 4:**
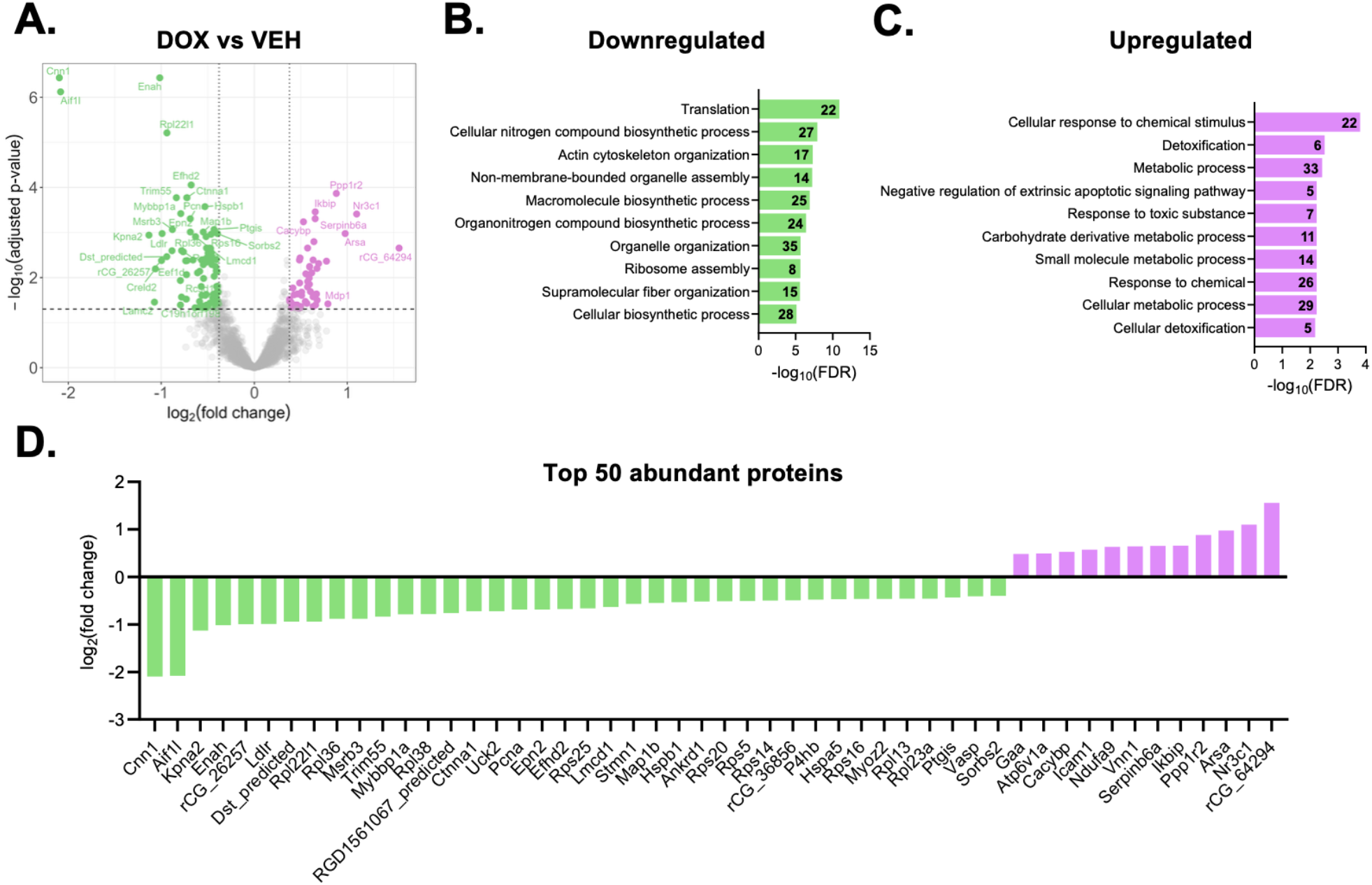
Cardiomyocyte proteome response to doxorubicin. Neonatal rat ventricular myocytes cultured in 10% newborn calf serum and treated with vehicle and 0.3 μM doxorubicin (Dox) for 24 hours. The limma package in R was used for differential abundance analysis of proteins, and Heatmapper used to generate heatmaps. a) Volcano plot of differentially abundant proteins indicated as purple (up) and green (down) with FDR <0.05 and |fold change| >1.3. 140 proteins were differentially abundant, including 51 upregulated and 89 downregulated. STRING analysis with Gene Ontology enrichment shows the top 10 Biological Processes enriched amongst the b) downregulated and c) upregulated proteins. d) Bar chart of the top 50 significantly differentially abundant proteins by FDR. FDR; false discovery rate.

#### Sex-specific response to DOX

Our limma model (which included treatment, sex and batch) did not detect an overall sex effect. However, there were clear differences in proteomic profiles between male and female DOX-treated samples (Fig 5A-B). Several proteins that have previously been shown to be differentially regulated by DOX in male and female mouse hearts were similarly regulated in our model (see annotated proteins on Fig 5C).^30,31^ This included proteins with known roles in DOX cardiac pathogenesis (cold inducible RNA binding protein, CIRBP;^32^ increased in female but not male hearts following DOX treatment^30^) and a biomarker for worsening heart failure (galectin-3, LGALS3;^33^ increased in male but not female hearts following DOX treatment^31^). In male myocytes, DOX treatment increased abundance of 7 proteins and decreased abundance of 22 proteins (Fig 5A). In female myocytes, DOX treatment increased abundance of 13 proteins and decreased abundance of 30 proteins (Fig 5B). Of these, only 13 proteins were common to both conditions, with 16 proteins significantly regulated by DOX in male myocytes only and 30 significantly regulated by DOX in female myocytes only (Fig 5D-G).

**Figure 5:**
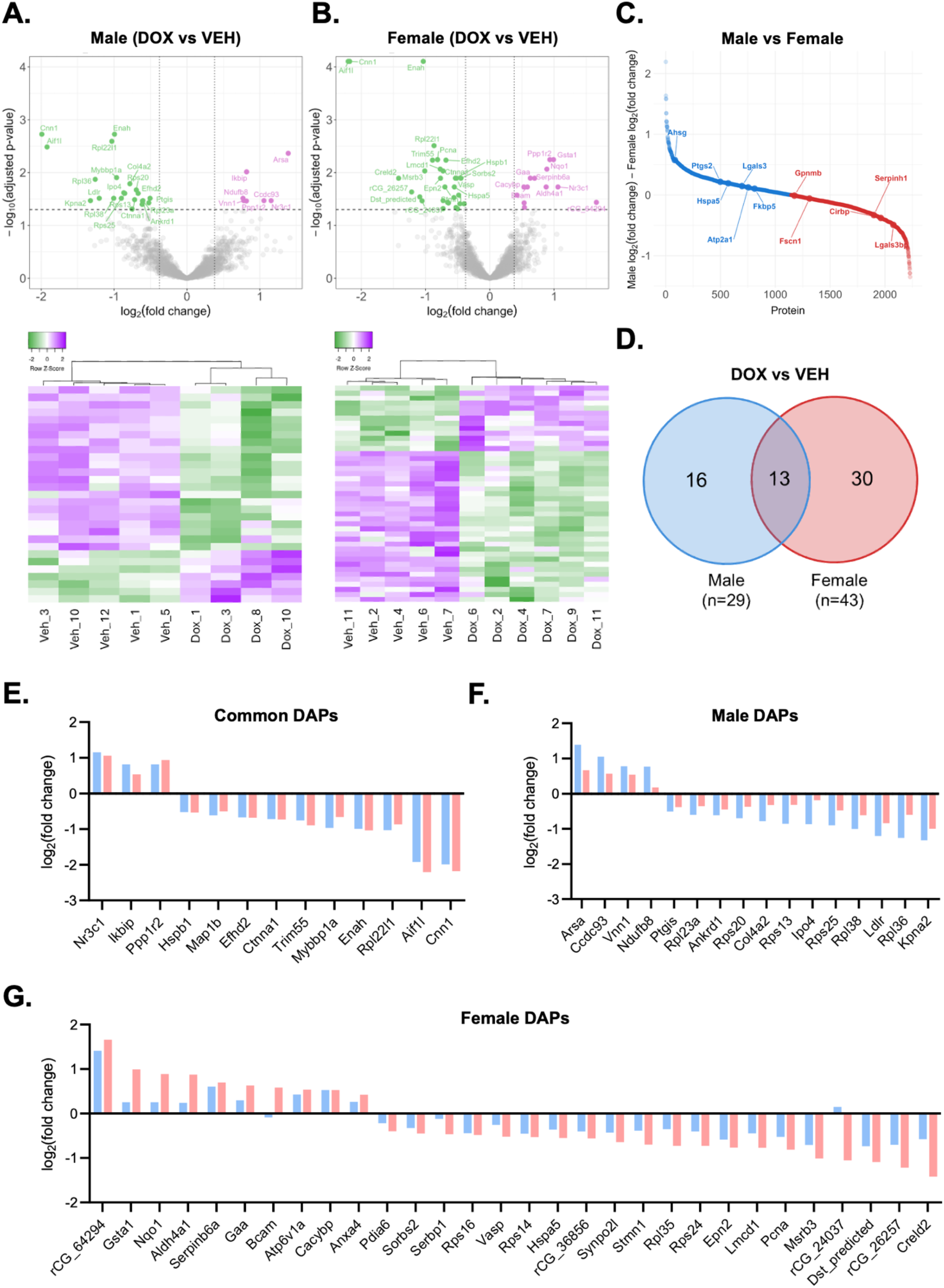
Sex differences in the cardiomyocyte proteome in response to doxorubicin. Neonatal rat ventricular myocytes cultured in 10% newborn calf serum and treated with vehicle and 0.3 μM doxorubicin (Dox) for 24 hours. The limma package in R was used for differential abundance analysis of proteins. Volcano plots of differentially abundant proteins indicated as purple (up) and green (down) with FDR <0.05 and |fold change| >1.3. Heatmaps generated in Heatmapper show Z-scores of normalised abundance values across samples, clustered hierarchically using Pearson distance and average linkage. a) Male analysis shows 29 proteins were differentially abundant, including 7 upregulated and 22 downregulated. b) Female analysis shows 43 proteins were differentially abundant, including 13 upregulated and 30 downregulated. c) Waterfall plot showing difference in DOX effect between males and females. d) Venn diagram comparing male and female differentially abundant proteins. Bar charts showing the log_2_FC for proteins that were significantly regulated by Dox in e) both sexes, f) male cardiomyocytes only, and g) female cardiomyocytes only.Blue, male; red, female; DAPs, differentially abundant proteins.

In summary, this dataset provides a valuable resource for exploring sex-specific cardiomyocyte protein changes in DOX-induced cardiotoxicity. In the analysis presented in this Data Descriptor, we filtered for proteins that were detected in 100% of samples to ensure high confidence in quantification. Allowing for missing values increases proteome coverage (Table 1) and may reveal additional candidates with sex-specific DOX treatment profiles. There is also potential value in examining proteins that were completely absent in one or more conditions (which were removed by our filtering threshold for the purposes of validating the dataset), as these may represent proteins that are most responsive to DOX and therefore of most interest biologically (Table 2, 3).

**Table 1:** Total protein counts with diPerent missing value thresholds.

| **Missing values** | **Protein count** |
| --- | --- |
| No missing values | 2229 |
| Up to 1 missing value in each group | 2905 |
| Up to 2 missing values in each group | 3518 |
| Up to 25% missing values (all samples) | 3379 |
| Up to 50% missing values (all samples) | 4170 |

**Table 2:** Proteins present only in vehicle- or in DOX-treated male cardiomyocytes.

| MALE | Gene name | Description | Samples (#) |
| --- | --- | --- | --- |
| **Present only in DOX** | Abcb1b | P-type phospholipid transporter | 4/4 |
|  | Tdrkh | Tudor and KH domain-containing protein | 4/4 |
|  | Cobll1_<br>predicted | Cobl-like 1 (Predicted), isoform CRA_a | 3/4 |
|  | Lars2_<br>predicted | leucine--tRNA ligase | 3/4 |
|  | Raph1_<br>predicted | Ras association (RalGDS/AF-6) and pleckstrin homology domains 1 (Predicted), isoform CRA_c | 3/4 |
|  | Prdm16 | [histone H3]-lysine(9) N-methyltransferase | 3/4 |
|  | Gas6 | Growth arrest-specific protein 6 | 3/4 |
|  | Mtrfr | Mitochondrial translation release factor in rescue | 3/4 |
|  | Sec3l1 | SEC3-like 1 (S. cerevisiae) (Fragment) | 3/4 |
|  | Steap4 | Ab1-219 | 3/4 |
|  | rCG_52086 | Elongator complex protein 3 | 3/4 |
| **Present only in VEH** | Ivns1abp | Influenza virus NS1A binding protein (Predicted), isoform CRA_b | 5/5 |
|  | Cog8_<br>predicted | Peptide deformylase | 5/5 |
|  | rCG_57706 | RCG57706 | 5/5 |
|  | rCG_43522 | DNA replication licensing factor MCM3 | 5/5 |
|  | Dnmt1 | DNA (cytosine-5)-methyltransferase | 5/5 |
|  | Rrm2 | Ribonucleoside-diphosphate reductase subunit M2 | 4/5 |
|  | Ncor1 | Nuclear receptor corepressor 1 | 4/5 |
|  | Lig3 | DNA ligase | 4/5 |
|  | Rsad1 | Radical S-adenosyl methionine domain-containing protein | 4/5 |
|  | Celf1 | CUG triplet repeat, RNA binding protein 1, isoform CRA_a | 4/5 |
|  | RGD1305256_<br>predicted | Similar to chromosome 20 open reading frame 75 (Predicted) (Fragment) | 4/5 |
|  | Desi1 | Desumoylating isopeptidase 1 | 4/5 |
|  | Trak1 | Similar to 106 kDa O-GlcNAc transferase-interacting protein (Predicted), isoform CRA_a | 4/5 |
|  | Mical2 | F-actin monooxygenase | 4/5 |
|  | Chst11 | Carbohydrate sulfotransferase | 4/5 |
|  | RGD1311900_<br>predicted | Major facilitator superfamily domain-containing protein 10 (Fragment) | 4/5 |
|  | Furin | Furin | 4/5 |
|  | Utp14a | RCG53178 | 4/5 |
|  | Ifngr1 | Interferon gamma receptor 1 | 4/5 |
|  | Ufsp2 | Ufm1-specific protease 2 | 4/5 |
|  | Ehbp1_<br>predicted | EH domain binding protein 1 (Predicted) | 4/5 |
|  | Samd4a | Protein Smaug homolog 1 | 4/5 |
|  | Cnot3_<br>predicted | CCR4-NOT transcription complex subunit 3 | 4/5 |
|  | Lig1 | DNA ligase | 4/5 |

**Table 3:** Proteins present only in vehicle- or in DOX-treated female cardiomyocytes.

| **FEMALE** | **Gene name** | **Description** | **Samples (#)** |
| --- | --- | --- | --- |
| **Present only in DOX** | Cdkn1a | Cyclin-dependent kinase inhibitor 1 | 6/6 |
|  | Dkk3 | Dickkopf-related protein 3 | 5/6 |
|  | Cdkn1c | Cyclin-dependent kinase inhibitor 1C | 5/6 |
|  | Pip4p2 | Phosphatidylinositol-4,5-bisphosphate 4-phosphatase | 5/6 |
|  | Slc27a3 | long-chain-fatty-acid--CoA ligase | 5/6 |
| **Present only in VEH** | Ajuba | Ajuba homolog (Xenopus laevis) | 5/5 |
|  | P4htm | Transmembrane prolyl 4-hydroxylase | 5/5 |
|  | Syf2 | Pre-mRNA-splicing factor SYF2 | 5/5 |
|  | Jcad | RCG45278, isoform CRA_a | 5/5 |
|  | Cdk1 | Cell division cycle 2 homolog A (S. pombe) | 5/5 |
|  | Ube2c | Ubiquitin-conjugating enzyme E2 C | 5/5 |
|  | Ascc1 | Activating signal cointegrator 1 complex subunit 1 | 4/5 |
|  | Ddx59 | Probable ATP-dependent RNA helicase DDX59 | 4/5 |
|  | Dyrk1a | Dual specificity tyrosine-phosphorylation-regulated kinase 1A | 4/5 |
|  | Nop14 | Similar to 2610033H07Rik protein (Predicted), isoform CRA_b | 4/5 |
|  | Ireb2 | Iron-responsive element-binding protein 2 | 4/5 |
|  | Rrm2 | Ribonucleoside-diphosphate reductase subunit M2 | 4/5 |
|  | Dnmt1 | DNA (cytosine-5)-methyltransferase | 4/5 |

## Data Availability

The mass spectrometry proteomics data have been deposited to the ProteomeXchange Consortium (http://proteomecentral.proteomexchange.org) via the PRIDE partner repository^22^ with the dataset identifier PXD076454.

## Code Availability

Input files and source code used for the bioinformatics analyses and data visualisation are provided at https://github.com/katelweeks/NRVM-DOX-proteomics.

## Author Contributions

S.D. – Conceptualisation, Methodology, Software, Validation, Formal analysis, Investigation, Data Curation, Writing – Original draft, Writing – Review & editing, Visualisation; M.L. – Software, Validation, Formal analysis, Resources, Data Curation, Writing – Review & editing; K.D. – Methodology, Resources, Writing – Review & editing; H.K. – Software, Validation, Formal analysis, Data Curation, Writing – Review & editing; E.J.H. – Writing – Review & editing, Supervision; B.L.P. – Resources, Visualisation, Writing – Review & editing; L.M.D.D. – Conceptualisation, Resources, Writing – Review & editing, Supervision; K.L.W. – Conceptualisation, Methodology, Investigation, Writing – Original draft, Writing – Review & editing, Supervision, Project administration.

